# Silvicultural practices and interannual variation shape ectomycorrhizal fungal diversity and community composition in an oak-hornbeam forest in northern Hungary

**DOI:** 10.64898/2026.04.28.721325

**Authors:** Kennedy Ododa, Péter Ódor, Bence Kovács, Flóra Tinya, Réka Aszalós, Carla Mota Leal, Adrienn Geiger, Anna Molnár, József Geml

## Abstract

Ectomycorrhizal (ECM) fungi are well-known for their crucial roles in forest health and productivity, yet their responses to various forest management practices are understudied, particularly in oak-dominated forests. The purpose of this study was to better understand the effects of silvicultural treatments on the diversity and community composition of ECM fungi in an oak-hornbeam forest in northern Hungary. We analyzed ITS2 rDNA metabarcoding data of soil-borne fungi to compare richness and community composition of ECM fungi among forest treatment types (clear-cutting, gap-cutting, preparation-cutting, tree retention in clear-cut areas, and control) and between sampling years (2020 and 2021). We found 268 ECM fungal genotypes, with the most diverse phylogenetic clades being /russula-lactarius (52), /tomentella-thelephora (47), /inocybe (40), /sebacina (27), and /cortinarius (20). We found significant compositional difference of ECM fungi among silvicultural treatments in both years, with some variations in richness. There were also small, but still significant compositional differences between the two years. Treatment effect was partly explained by altered environmental variables, such as relative humidity and soil temperature. These results highlight the importance of forest structure and the abiotic environment in driving community dynamics of plant-symbiotic fungi, with potential implications for forest health and productivity.

## Introduction

The interactions between forest tree composition, stand structure and forest microbiome have important ramifications for forest biomass production as well as ecosystem functioning and ecosystem services, but our understanding of these interactions and how they are influenced by silvicultural practices and local environmental variables. In temperate forests, ectomycorrhizal (ECM) fungi constitute one of the most ecologically and economically important components of the forest soil microbiome, because ECM fungi are critical symbiotic partners of most forest trees, including oaks and beeches (*Fagaceae*), hornbeams, birches and alders (*Betulaceae*) as well as pines, spruces and firs (Pinaceae) (Berch *et al*., 2023; Molina and Horton 2015). These fungi facilitate the host’s acquisition of water and essential nutrients, notably nitrogen and phosphorus, thereby supporting tree growth and health. In addition, ECM fungi enhance host resilience against abiotic stressors such as drought and soil nutrient limitation, while contributing significantly to key ecosystem processes including soil carbon sequestration and nutrient cycling (Ekholm *et al*., 2023; Kuyper & Suz, 2023; Shan *et al*., 2025). In the Pannonian biogeographic region, where our study site is located, most forests are dominated by various oak (*Quercus*) species, with the landscape-level spatial distribution of the different forest types strongly influenced by topography-driven mmesoclimatic and edaphic conditions (Geml 2019). Previous studies have identified abiotic drivers such as soil pH, moisture availability, and topography-driven differences in mesoclimate as important influences on ECM fungal community patterns at landscape scales in Pannonian forests (Geml 2019; Geml *et al*., 2022) and in other temperate forests alike (Crowther *et al*., 2014; Castaño et al. 2018; Mandolini *et al*., 2024).

Silvicultural practices impose direct modifications of the forest stand structure that affects microclimate and soil conditions, e.g., soil chemistry, light penetration, moisture regimes, and the availability of compatible host trees (Kovács *et al*. 2020; Rianhard *et al*., 2025a; Tomao *et al*., 2020). Intensive interventions, such as clear-cutting, frequently result in marked reductions in biodiversity in general (Ekholm *et al*., 2023; Tinya *et al*., 2023) and in ECM fungi in particular, with shifts in community composition often persisting for decades (Lunde *et al*., 2025). In contrast, management techniques with less changes in stand structure and microclimate, including gap-cutting, retention forestry, and low-impact thinning, have shown to have less impact on ECM fungal diversity (Rosenvald & Lõhmus, 2008; Sterkenburg *et al*., 2019). However, these findings are not universally consistent: while some studies report resilience and functional redundancy of ECM communities under moderate disturbance (Wilgan & Leski, 2022), others document lasting alterations in the composition and functionality of ECM fungal communities that are critical for nutrient uptake (Tomao *et al*., 2020). Furthermore, most publications on the effects of forest management on ECM fungi in Europe focused on beech and conifer forests, while similar studies are scarce for oak forests, particularly in Central Europe. Currently, there is an important gap in our knowledge regarding the interactions of forest management and ECM fungal communities that crucial for forest health and productivity.

In this paper, we generated and analyzed soil DNA metabarcoding data to assess the compositional dynamics of ECM fungal communities under various silvicultural practices. We built on the research infrastructure of the long-term the Pilis Forestry Systems Experiment in northern Hungary, a multi-taxon forest ecological study to characterize the effects of various silvicultural practices, representing components of rotation and continuous-cover forestry systems, on biotic communities and forest regeneration (https://piliskiserlet.ecolres.hu/en). We investigated how ECM fungal communities respond to five silvicultural treatments: clear-cutting, gap-cutting, preparation-cutting (thinning), tree retention in clear-cut areas, and unmanaged control. These treatments represent a gradient of disturbance intensity and host tree removal. Using soil DNA sequence data generated from samples collected in 2020 and 2021, combined with environmental data on microclimate and edaphic factors and understory vegetation, this research assesses spatial and temporal variation in ECM fungal communities six and seven years after the implementaztion of the treatments during the winter of 2014-2015.

Our analysis focuses on three main objectives: (1) evaluating how ECM fungal richness vary among silvicultural treatments; (2) evaluating the effects of silvicultural practices of various intensity on the composition of ECM fungal communities; and (3) assessing the interannual variability of ECM fungal communities at the same sites. More specifically, we expected to see differences in the diversity ECM fungal communities among treatments, particularly between control and clear-cut plots, representing the least and most intense disturbance, respectively (Hypothesis I). Based on niche theory, we expected to see significant compositional differences of ECM fungi among silvicultural treatments, because of the altered environmental conditions (Hypothesis II). Finally, we hypothesized that there would be little interannual variation in ECM fungal communities, because many ECM fungi have extensive vegetative mycelial structures that often persist for years (Hypothesis III).

## Materials and methods

### Study area and experimental design

The study was conducted in the Pilis Forestry Systems Experiment (47°40 N, 18°54 E), located the Pilis Mountains at the northeastern part of the Transdanubian Range in northern Hungary (Fig. 1; Kovács *et al*., 2020). The studied stand is a typical mature (80 years old) Pannonian oak-hornbeam forest (Natura 2000 code: 91G0) managed under the traditional shelterwood (rotational) forestry system. Sessile oak (*Quercus petraea*) dominates the upper canopy layer, with hornbeam (*Carpinus betulus)* forming a secondary canopy layer. Subordinate species include *Quercus cerris, Prunus avium, Fagus sylvatica*, and *Fraxinus ornus*. There is a sparse ground layer of general and mesic forest species, with very little shrub layer. *Melica uniflora* and *Carex pilosa* are the dominant understory species. Plots are located at 370–450 meters above sea level on a gentle north-facing slope. The mean annual temperature is between 9.0 and 9.5 °C, and the mean annual precipitation is between 600 and 650 mm (Dövényi, 2010). The most prevalent soil type is lessivage brown forest soil (luvisol), which is slightly acidic, with pH of the top 20 cm layer between 4.2 and 5.3 (Fekete & Tóth, 2010; Kovács *et al*., 2018). The bedrock is limestone and sandstone with loess.

**Figure 1.**
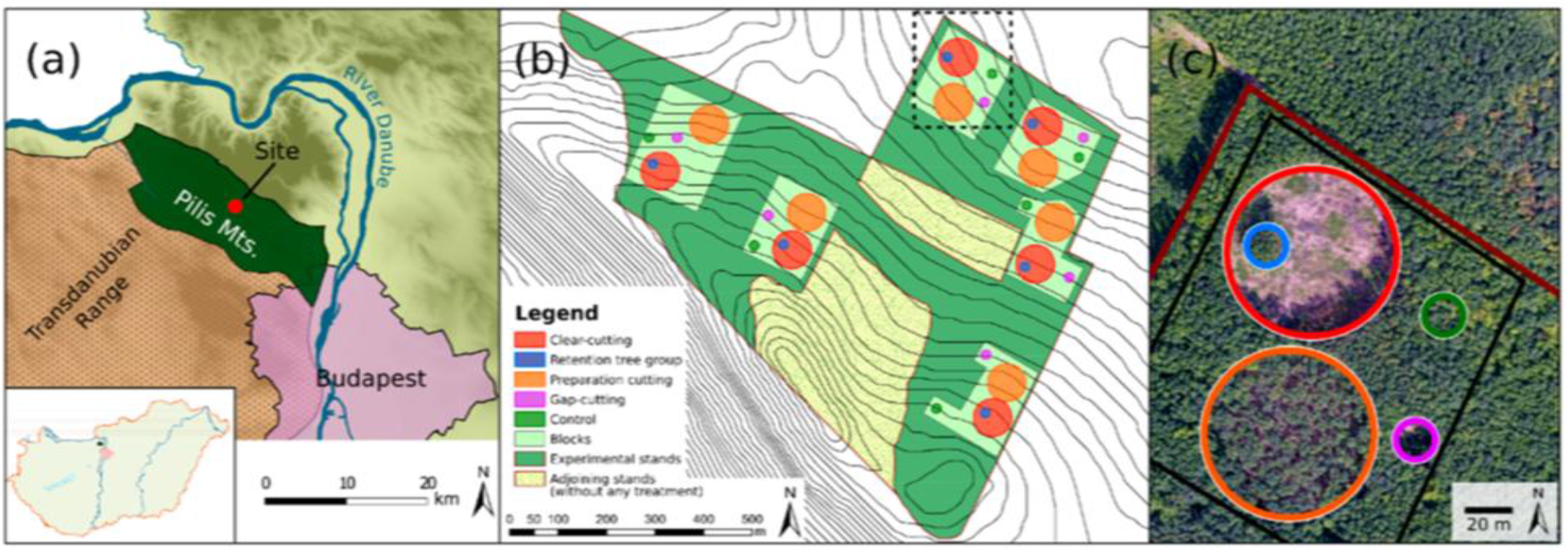
(a) Location of the experimental area in Hungary. (b) Map of the experimental area with the six blocks comprising five plots. (c) A drone image of one block showing the silvicultural treatments: clear-cutting (red) with retention tree group (blue), gap-cutting (magenta), preparation-cutting (orange), and unaltered control (green). The drone image was provided by Dr. Viktor Tóth of the Ecological Research Centre in collaboration with Pilis Park Forest Company.

The experimental design included four silvicultural treatments: clear-cutting (CC), retention tree group (R), preparation-cutting (P), gap-cutting (G), and the control (C) in six blocks using completely randomized block design, resulting in 30 plots (Figure 1). Specifics of the treatments were as follows: The treatments were as follows: control (C): representing a closed-canopy stand without intervention; clear-cutting (CC): a circular area, where all trees with DBH ≥ 5 cm and/or height ≥ 2 m were removed (diameter 80 m, area 0.5 ha); gap-cutting (G): an artificial circular gap with a gap diameter to intact canopy height ratio of approximately 1:1 (diameter 20 m, area 0.03 ha); preparation cutting (P): uniform partial cutting within a circle of 80-meter diameter, where 30% of the total basal area of the dominant canopy layer and the entire secondary canopy layer were removed in a spatially even arrangement; and retention tree group (R): within the clear-cuts, 8–12 dominant tree and shrub individuals were retained as a circular patch of 0.03 hectares (20 m diameter). The stages of P, R, and CC are typical of the rotational forestry system, while G is frequently used within the framework of the continuous cover forestry system. The treatments were implemented during the winter of 2014-2015. Soil samples were collected in the first week of October in 2020 and 2021.

### Sampling and molecular work

In each plot, eight top soil cores, 10 cm deep, were taken at least 2 m apart from underneath the litter layer, using a cylindrical soil corer (5 cm in diameter), and were mixed to provide a composite sample per plot. Genomic DNA was extracted from 0.5 ml of soil from each composite sample using the NucleoSpinR soil kit (Macherey-Nagel Gmbh & Co., Düren, Germany), following the manufacturer’s instructions. The DNA sequencing and PCR procedures were followed exactly as outlined in (Geml *et al*., 2014). In summary, the ITS2 region (about 250 bp) of the nuclear ribosomal rDNA repeat was amplified using fungal-specific primers fITS7 (Ihrmark *et al*., 2012) and ITS4 (White *et al*., 1990) with Illumina adapters under the following PCR conditions: one cycle of 95 °C for 5 min, followed by 37 cycles of 95 °C for 20 s, 56 °C for 30 s, and 72 °C for 1.5 min, culminating in a single cycle of 72 °C for 7 min, with negative controls. Forward and reverse primers were tagged with a combination of two distinct eight-nucleotide labels, resulting in a unique combination for each sample, in order to link the sequences to the sample source. The sequencing the company used default positive and negative controls to generate 250-bp paired-end reads from the amplicon libraries, which were then sequenced using Illumina MiSeq at Eurofins BIOMI (Gödöllő, Hungary) after being normalized for DNA concentration.

### Microclimate data

We have been gathering daily microclimate data in all 30 plots since May, 2020 to present with hourly measurements. For statistical analyses in the manuscript, we used microclimate data from May through September in 2020 and 2021, spanning 5 months of the vegetative period immeditely before the fall sampling each year. Two categories of aggregated data were established: absolute values and relative values. All treatment plot direct measurements were included in the absolute data, whereas values computed as block-by-block variations from the control plots formed the relative data. The measured variables included air temperature (°C) measured at heights of 130 cm, 15 cm, and on the ground surface (T130, T15, and T0), relative humidity (RH in %) measured at 130 cm above ground, vapor pressure deficit (VPD in kPa), calculated from T130 and RH values using the plantecophys::RHtoVPD method, soil temperature (°C) measured 8 cm below the ground (Tsoil), volumetric water content (VWC) calculated from TDT counts using specific soil class coefficients. Voltcraft DL-210TH loggers were used to measure RH and T130), while TOMST TMS-4 loggers recorded Tsoil, T0, and T15, and calculated VWC). To depict the microclimate conditions, daily data for each variable were summarised using averages, maxima, minima, inter-quartile ranges, and percentiles (5% and 95%).

### Bioinformatic and statistical analyses

Unless otherwise noted, all bioinformatic and statistical analyses were carried out in R v. 4.3.2 (R Development Core Team, 2021). The raw DNA sequences were analyzed using the dada2 package (Callahan *et al*., 2016) in R (R Core Team, 2020). Based on the quality score profiles, forward reads were truncated to 250 bp and reverse reads to 200 bp. Furthermore, a quality-filtering step was applied, with the maximum expected errors (maxEE) allowed in a read set to 2. The filtered reads underwent denoising, after which the two-directional reads were merged and clustered into amplicon sequence variants (ASVs), which were subsequently filtered for chimeric sequences. Because of the large per-sample sequencing depth of 306,216 high-quality reads, only ASVs with at least 10 sequences in a given sample were deemed “present” in that sample to reduce false presences. Furthermore, to prevent artifactual ASVs, ASVs found in a single sample were excluded from further analyses. Taxonomic classifications of the sequences were determined with USEARCH v. 11 (Edgar, 2010), using the version April 4, 2024 of the UNITE database of reference sequences (Abarenkov *et al*., 2024). Only ASVs with at least a 80% sequence similarity to a fungal reference sequences were retained for further analyses. We assigned ASVs identified to at least the genus level to functional groups based on the curated reference database of FungalTraits version 1.2 (Põlme *et al*., 2020). For this paper, we selected all ECM fungal ASVs for in-depth analyses and these were assigned to phylogenetic lineages of ECM fungi *sensu* Tedersoo and Smith (2013) based on the assignation of the matching SHs in UNITE. Molecular data has been deposited deposited at DDBJ/EMBL/GenBank in a Targeted Locus Study project under the accession KJLF00000000. The version described in this paper is the first version, KJLF01000000.

Alpha diversity, measured as ASV richness, was compared among treatment types and years using one-way ANOVA after checking assumptions of normality (Shapiro-Wilk test) and homogeneity of variance (Levene’s test). Post hoc multiple comparisons were conducted with Tukey’s HSD test. For specific pairwise comparisons, independent t-tests were applied. Where assumptions were not met, non-parametric tests were used instead. Beta diversity was analyzed using permutational multivariate analysis of variance (PERMANOVA) with the *adonis2* function from the *vegan* package (Oksanen et al. 2015). The analysis used a Hellinger-transformed abundance matrix and Bray–Curtis distance to test differences in community composition among treatments and years. Non-metric multidimensional scaling (NMDS) was performed on the same transformed data to visualize patterns of compositional differences among samples using the *metaNMDS* function. Stress values from NMDS were used to evaluate the quality of the ordination. Indicator species analysis was conducted with the *multipatt* function in the *indicspecies* package (De Cáceres et al. 2012) to identify taxa associated with specific treatments and years. The statistical significance of indicator taxa was determined by permutation tests. Abundances of indicator species were correlated with environmental variables, including soil moisture, relative humidity, soil temperature, understory vegetation, and proximity to ECM host trees. Pearson or Spearman correlation tests were applied depending on data distribution. Graphs of ASV richness and relative abundances were produced using the *ggplot2* package (Wickham 2016), including boxplots and scatterplots with confidence intervals. Effect sizes such as Cohen’s d were calculated alongside significance tests to quantify the magnitude of differences.

## Results

Of the 1696 fungal ASVs detected, 268 represented ECM fungi. The most diverse phylogenetic clades of ECM fungi in the samples were /russula-lactarius (52 genotypes), /tomentella-thelephora (47), /inocybe (40), /sebacina (27), and /cortinarius (20). Results indicated a significant treatment and year effect on total ECM richness, *F*(9, 49) = 4.06, *p* < 0.001, partial η^2^ = 0.43 (Figure 2). The 2021 samples from clear-cut plots (CC21) showed the lowest richness, significantly less than richness values observed in the 2021samples from the control (C21) and preparation (P21) plots. Other treatment × year combinations showed intermediate richness, with no significant differences (Figure 2).

**Figure 2.**
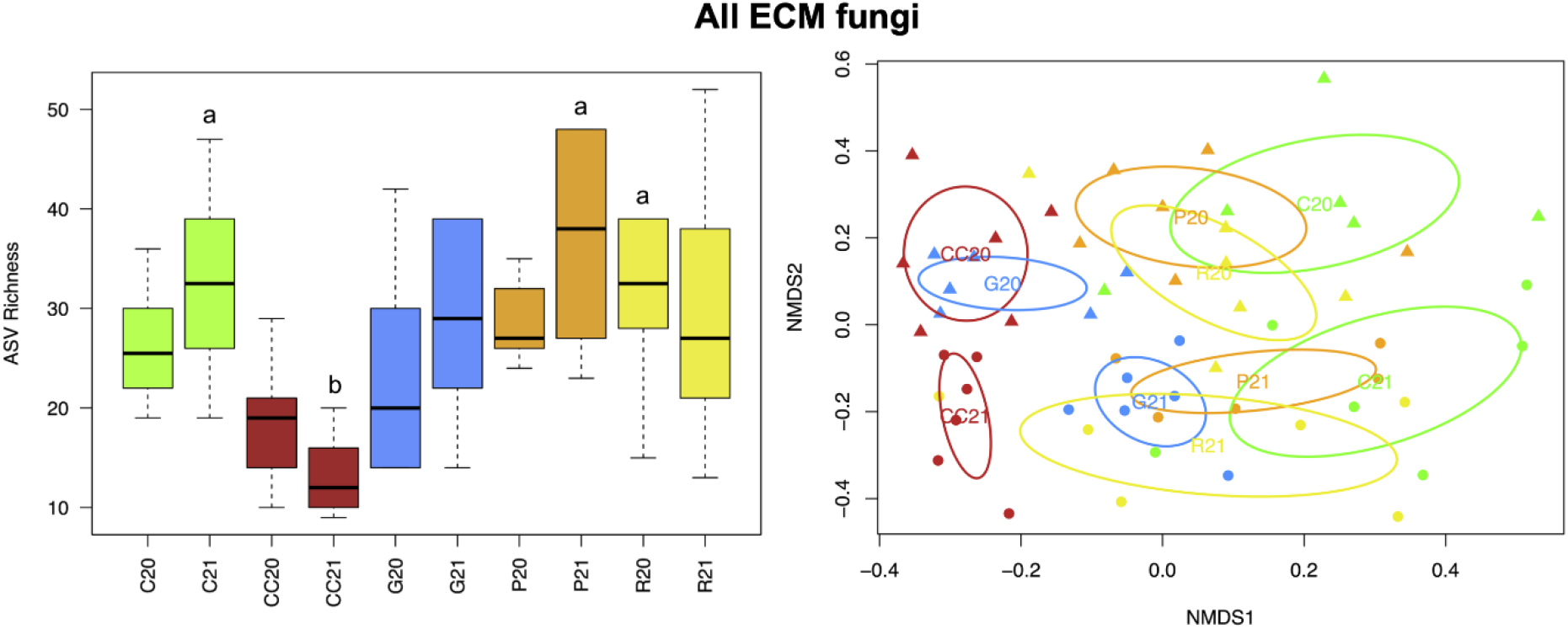
All ECM fungal richness among treatment × year combinations, with significant differences are indicated by letters (left). Non-metric multidimensional scaling (NMDS) ordination plot (right) showing the compositional differences of ECM fungal communities among silvicultural treatments based on Bray-Curtis distances of Hellinger-transformed abundance data (stress = 0.1707017). Abbreviations for silvicultural treatments are C: unaltered control (green), CC: clear-cutting (red), G: gap-cutting (blue), P: preparation-cutting (orange), and R: retention tree group (yellow).

Several phylogenetic clades ECM fungi showed differences in richness among silvicultural treatments and occasionally between sampling years, mostly with higher richness in plots with no or small disturbance (C, P, and R) than in CC and, to a lesser extent, G plots, although the differences were not always significant (Figure 3). Richness in /amanita differed significantly across treatment-year combinations, F(9, 49) = 2.15, p = 0.042, η^2^ = 0.28, with the highest values observed in the control plots (C20 and C21) and consistently lower richness in most other treatments. In /cenococcum, richness varied more markedly, F(9, 49) = 5.11, p < 0.001, η^2^ = 0.48; the C21 plot recorded the highest richness, while clear-cut treatments (CC20 and CC21) had the lowest values. Gap-cutting treatments (G20 and G21) exhibited intermediate richness, with other treatments showing moderate values. Richness in /inocybe ASVs also differed significantly, F(9, 49) = 3.07, p = 0.005, η^2^ = 0.36, though most pairwise comparisons were not significant except for contrasts such as between R20 and CC21. Richness of the /russula-lactarius clade varied significantly across treatment × year combinations, F(9, 49) = 7.07, p < 0.001, η^2^ = 0.57, with the highest richness found in the control plots and P21, and the lowest in clear-cut plots (CC20 and CC21). Other treatments, including retention, showed moderate richness levels. The /tomentella-thelephora also showed a significant treatment effect, F(9, 49) = 2.77, p = 0.011, η^2^ = 0.34, with the lowest richness in CC21 and higher values in P21 and R20; several treatment-year combinations grouped at intermediate levels (Figure 3).

**Figure 3.**
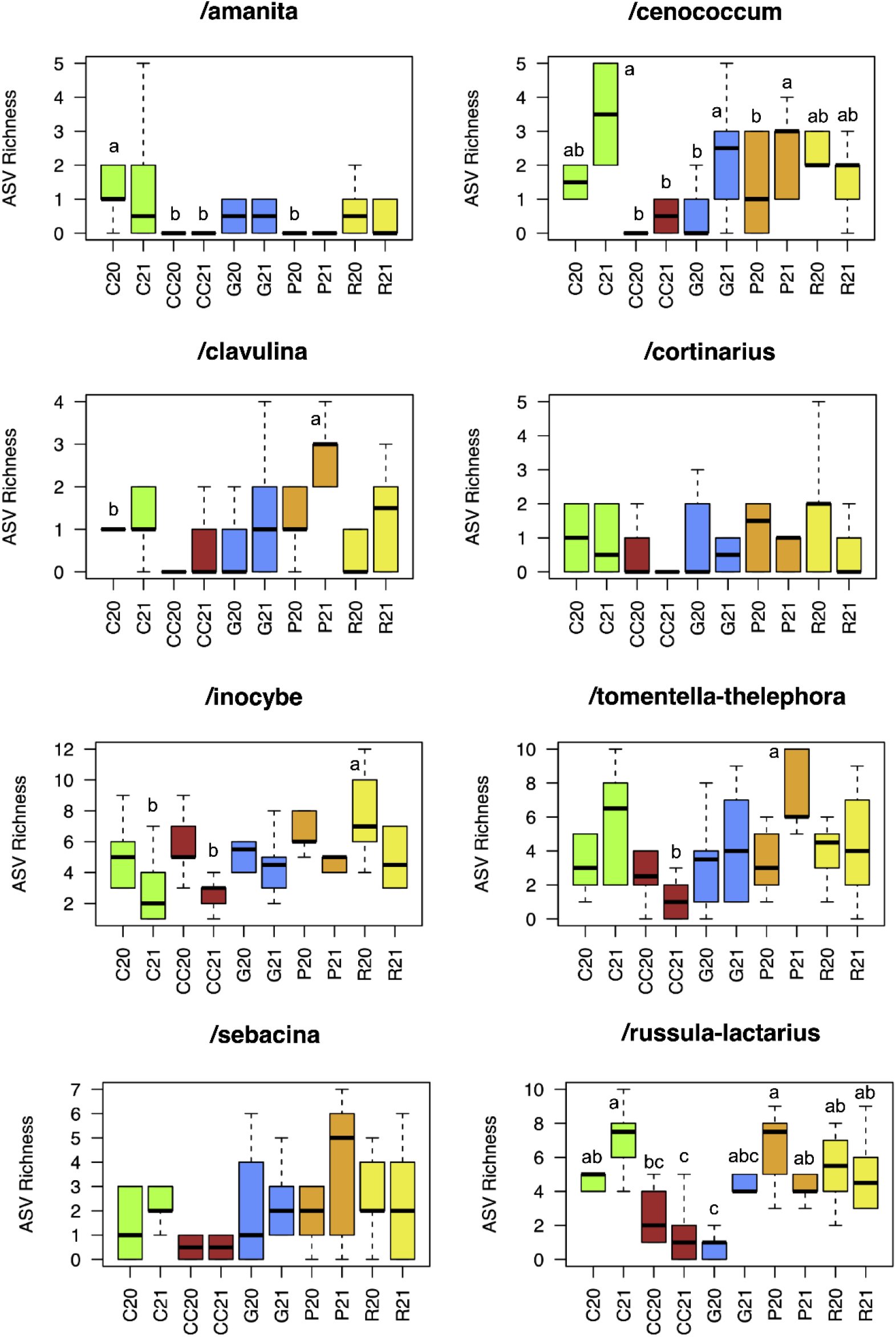
ASV richness of the five most diverse ECM fungal genera among treatment × year combinations. Significant differences are indicated by letters. Abbreviations for silvicultural treatments follow those in Figure 2.

Compositional differences among silvicultural treatments were significant in both sampling years with respect to the entire ECM fungal community as well as the phylogenetic lineages tested (Figures 2 and 4). Generally, the first axis correlated with treatments, highlighting the level of intensity of the treatment, with CC and G plots being the most different from the unaltered controls (C), while the second axis showed the difference between the two sampling years (Figure 2).

**Figure 4.**
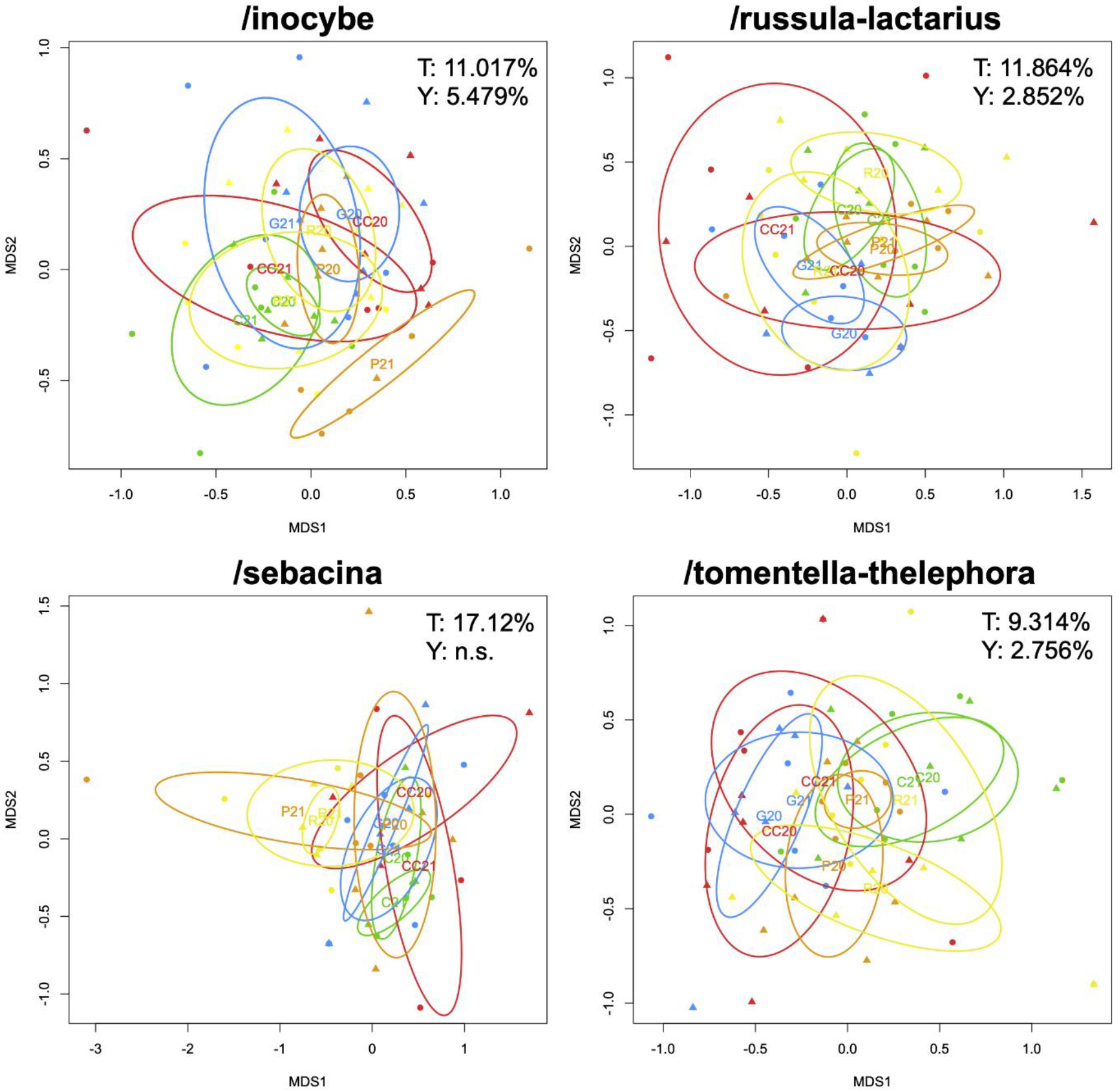
Non-metric multidimensional scaling (NMDS) ordination plots showing the compositional differences of the four dominant ECM fungal lineages among silvicultural treatments based on Bray-Curtis distances of Hellinger-transformed abundance data. Compositional variance (in %) significantly explained by treatments (T) and years (Y) are indicated. Abbreviations for silvicultural treatments follow those in Figure 2.

The PERMANOVA analyses revealed that the composition of ECM fungal communities was primarily influenced by silvicultural treatment and related differences in microclimatic variables, while internnual differences were smaller and sometimes insignificant (Table 1). No significant interaction was found between treatment and year. Of the edaphic variables tested, only soil pH and organic matter content showed significant correlations with community turnover and only in /sebacina and /russula-lactarius, respectively. Among biotic variables, understory plant species richness and cover correlated significantly with fungal community composition in all ECM fungi, /inocybe, and /sebacina. Of the dominant tree species, only the abundance of sessile oak showed significant correlation: with the total ECM community, /russula-latarius, and /tomentella-thelephora. With respect to microclimatic variables, mean and interquartile range air temperature measured at various heights (0, 15, and 130 cm) and interquartile range of relative humidity, vapour pressure deficit, and soil water content explained a significant proportion of the compositional variance in all analyses (Table 1).

**Table 1.** Correlations of categorical variables silvicultural treatment and year, and continuous variables corresponding to soil chemistry, vegetation, and microclimate with compositional differences in all ECM fungal communities and in the dominant lineages, based on PERMANOVA analyses, where experimental blocks were treated as strata. Values indicating significant correlations are in bold, with continuous variables with significant contributions to the combined model accounting for correlations among variables are in italics. Abbreviations: OM – soil organic matter, N – soil nitrogen content, C – soil carbon content, AlP_2_O_5_ – phosphorus concent in the form of AlP_2_O_5_, AlK_2_O – potassium content in the form of AlK_2_O, Cover – understory plant cover, SpNo – understory plant species richness, Carbet – *Carpinus betulus* abundance, Fagsyl – *Fagus sylvatica* abundance, Quecer – *Quercus cerris* abundance, Quepet – *Quercus petraea* abundance, RH – relative humidity, T130 – air temperature at breast height (130 cm), T15 – air temperature at 15 cm height, T0 – soil surface temperature, Tsoil – soil temperature at 8 cm depth, VPD – Vapour Pressure Deficit, VWC – volumetric water content, iqr – interquartile range.

| Variables | All ECM fungi |  | /incocybe |  | /russula-lactarius |  | /sebacina |  | /tomentella-thelephora |  |
| --- | --- | --- | --- | --- | --- | --- | --- | --- | --- | --- |
| | $R^2$ | $p$ | $R^2$ | $p$ | $R^2$ | $p$ | $R^2$ | $p$ | $R^2$ | $p$ |
| Treatment | **0,12673** | **0,001** | **0,11017** | **0,004** | **0,11864** | **0,001** | **0,17120** | **0,001** | **0,09314** | **0,005** |
| Year | **0,06245** | **0,001** | **0,05479** | **0,001** | **0,02852** | **0,044** | 0,01625 | 0,773 | **0,02756** | **0,039** |
| Soil pH | 0,01644 | 0,543 | 0,01272 | 0,754 | 0,01719 | 0,439 | **0,04570** | **0,013** | 0,01946 | 0,269 |
| OM | 0,01963 | 0,211 | 0,02538 | 0,058 | **0,02870** | **0,048** | 0,02983 | 0,236 | 0,01436 | 0,575 |
| N | 0,01877 | 0,302 | 0,02000 | 0,211 | 0,02313 | 0,176 | 0,03044 | 0,246 | 0,01540 | 0,487 |
| C | 0,01883 | 0,323 | 0,02408 | 0,087 | 0,02366 | 0,141 | 0,02726 | 0,403 | 0,01560 | 0,541 |
| $\text{AlP}_2\text{O}_5$ | 0,01690 | 0,496 | 0,01542 | 0,571 | 0,01888 | 0,356 | 0,02896 | 0,254 | 0,01455 | 0,625 |
| $\text{AlK}_2\text{O}$ | 0,02164 | 0,096 | 0,01927 | 0,343 | 0,03382 | 0,023 | 0,04304 | 0,075 | 0,02167 | 0,204 |
| Cover | **0,03461** | **0,001** | **0,04833** | **0,002** | 0,01918 | 0,338 | **0,04115** | **0,037** | 0,02608 | 0,053 |
| SpNo | 0,02065 | 0,093 | **0,02972** | **0,012** | 0,02222 | 0,125 | **0,06317** | **0,001** | 0,02355 | 0,133 |
| Carbet | 0,01821 | 0,382 | 0,02936 | 0,092 | 0,01567 | 0,602 | 0,02351 | 0,511 | 0,02049 | 0,315 |
| Fagsyl | 0,01466 | 0,801 | 0,01456 | 0,734 | 0,01583 | 0,498 | 0,03584 | 0,088 | 0,01596 | 0,579 |
| Quecer | 0,01870 | 0,269 | 0,02140 | 0,398 | 0,01302 | 0,812 | 0,02235 | 0,756 | 0,01446 | 0,795 |
| Quepet | **0,02857** | **0,002** | 0,02978 | 0,211 | **0,03519** | **0,002** | 0,02222 | 0,893 | **0,03320** | **0,012** |
| T130_mean | **0,02405** | **0,026** | 0,02596 | 0,069 | 0,02661 | 0,078 | 0,01922 | 0,575 | **0,02731** | **0,028** |
| RH_mean | 0,01730 | 0,425 | 0,01251 | 0,792 | 0,01681 | 0,522 | 0,02563 | 0,373 | 0,02235 | 0,118 |
| VPD_mean | 0,01704 | 0,443 | 0,01064 | 0,856 | 0,02326 | 0,179 | 0,02369 | 0,438 | 0,02146 | 0,216 |
| T15_mean | **0,04615** | **0,001** | 0,02440 | 0,093 | **0,03723** | **0,006** | 0,02693 | 0,341 | **0,03013** | **0,015** |
| T0_mean | **0,03586** | **0,001** | 0,02524 | 0,062 | **0,03720** | **0,013** | 0,02939 | 0,207 | 0,03136 | 0,058 |
| Tsoil_mean | 0,02285 | 0,115 | 0,02699 | 0,134 | 0,02813 | 0,057 | 0,02864 | 0,528 | **0,03204** | **0,045** |
| VWC_mean | 0,01997 | 0,261 | 0,01555 | 0,781 | **0,03460** | **0,011** | 0,02678 | 0,624 | 0,01258 | 0,764 |
| T130_iqr | **0,05084** | **0,001** | **0,03133** | **0,011** | **0,03254** | **0,014** | 0,03085 | 0,185 | 0,02344 | 0,084 |
| RH_iqr | **0,04987** | **0,001** | **0,03299** | **0,011** | **0,03165** | **0,018** | 0,03516 | 0,099 | **0,02624** | **0,022** |
| VPD_iqr | **0,05105** | **0,001** | 0,03331 | 0,007 | **0,03240** | **0,014** | 0,03016 | 0,214 | 0,02309 | 0,083 |
| T15_iqr | **0,05336** | **0,001** | **0,04542** | **0,005** | **0,02967** | **0,03** | 0,02714 | 0,273 | 0,02344 | 0,111 |
| T0_iqr | **0,03971** | **0,001** | **0,03707** | **0,009** | 0,02190 | 0,176 | **0,04595** | **0,013** | 0,02532 | 0,083 |
| Tsoil_iqr | **0,03871** | **0,001** | **0,03856** | **0,003** | 0,02368 | 0,121 | 0,02919 | 0,186 | **0,03200** | **0,006** |
| VWC_iqr | **0,03924** | **0,001** | **0,04648** | **0,001** | 0,01799 | 0,391 | **0,04835** | **0,012** | **0,02990** | **0,016** |

Silvicultural treatments showed significant effects on the composition of all four dominant ECM fungal lineages, while compositional differences between years were much smaller, although still significant, except in /sebacina (Figure 4). The indicator species analysis identified distinct ASVs significantly associated with specific silvicultural treatments. Preparation-cutting and control plots had the most indicators, eight and five, respectively (Table 2).

**Table 2.** Indicator ASVs characteristically associated with forestry treatments, with corresponding *p*-values and with ITS2 rDNA sequence similarity (%), taxonomic assignment, and Species Hypothesis (SH) number of the most similar reference sequence in the UNITE database.

| ASV | Treatment | *p* | % similarity | Matching taxon | Species Hypothesis |
| --- | --- | --- | --- | --- | --- |
| ASV00269 | C | 0.0244 | 98.2 | *Amanita citrina* | SH1192419.09FU |
| ASV01182 | C | 0.0365 | 100 | *Byssocorticium atrovirens* | SH1187877.09FU |
| ASV00256 | C | 0.0329 | 100 | *Inocybe asterospora* | SH0886565.09FU |
| ASV00277 | C | 0.0311 | 97.9 | *Russula grata* | SH1206843.09FU |
| ASV00285 | C | 0.0044 | 97.1 | *Tomentella* sp. | SH1140071.09FU |
| ASV00383 | CC | 0.0213 | 97.1 | *Laccaria trichodermophora* | SH1173939.09FU |
| ASV00211 | G | 0.0278 | 92.7 | *Hymenogaster* sp. | SH1178886.09FU |
| ASV00245 | G | 0.0089 | 95.7 | *Hymenogaster* sp. | SH1179477.09FU |
| ASV00627 | G | 0.0195 | 100 | *Sebacina incrustans* | SH1295699.09FU |
| ASV00770 | P | 0.004 | 100 | *Cortinarius roseocastaneus* | SH1282078.09FU |
| ASV01143 | P | 0.0344 | 99.7 | *Inocybe minima* | SH1212493.09FU |
| ASV00179 | P | 0.0311 | 99.6 | *Inocybe* sp. | SH0186387.09FU |
| ASV00037 | P | 0.0056 | 97.2 | *Russula byssina* | SH1176998.09FU |
| ASV00162 | P | 0.0213 | 97 | *Russula nigricans* | SH1195370.09FU |
| ASV00059 | P | 0.0127 | 99.3 | *Sebacina* sp. | SH1232010.09FU |
| ASV01801 | P | 0.0291 | 99.7 | *Sebacina* sp. | SH1310638.09FU |
| ASV02626 | P | 0.0295 | 97.8 | *Tomentella* sp. | SH0926607.09FU |
| ASV01647 | R | 0.0322 | 97.7 | *Cortinarius olivaceoluteus* | SH1303684.09FU |
| ASV00115 | R | 0.0315 | 100 | *Sebacina* sp. | SH2076008.09FU |

We observed significant differences in understory plant cover and plant species richness among silvicultural treatments, with cover being significantly higher in all treatments compared to the control and the number of species being highest in the P and R plots (Figure 5). Similarly, microclimate in the treatment plots differed significantly from the control in most measured variables. In general, while mean air temperature measured at breast height (130 cm) did not differ significantly from the control, mean air temperature measurements at lower heights as well as soil temperature were significantly lower in treatments with tall and dense understory layer, i.e. CC and G, than teperatures measured in the control plots. Daily fluctiations in all measured microclimatic variables, as expressed by the inter-quartile-range (IQR), were significantly greater in the treatments than in the control (Figure 5).

**Figure 5.**
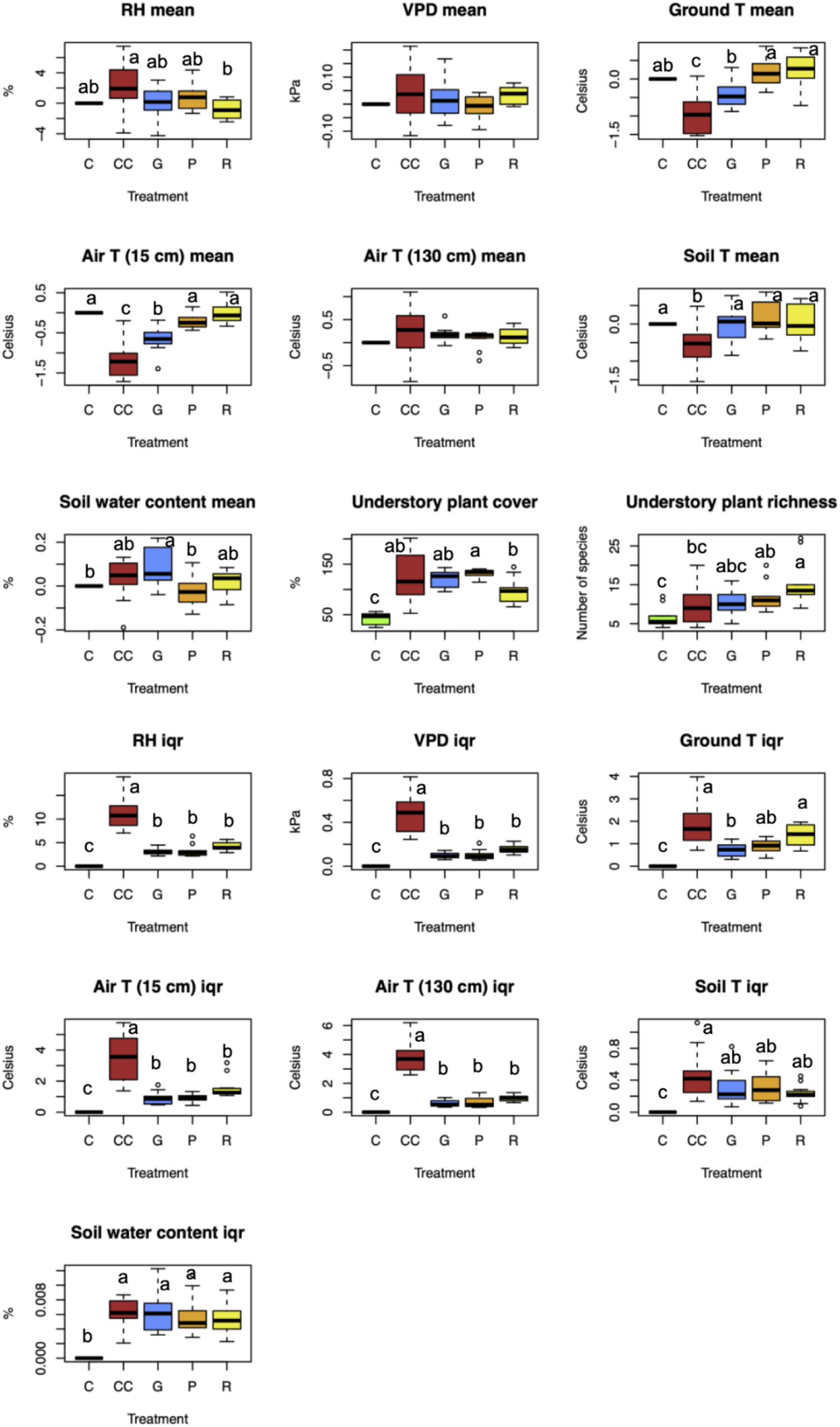
Changes in microclimatic variables and in understory plant cover and number of species as a result of the silvicultural treatments. In abiotic variables, the changes are expressed as differences from the control calculated at block level. Significant differences are indicated by letters. Abbreviations for silvicultural treatments follow those in Figure 2.

## Discussion

This study, integrating soil DNA metabarcoding and environmental data collected in plots subjected to experimental silvicultural treatments in a long-term setup, provides novel insights into the effects of forestry practices and the resulting environmental changes on the diversity and composition of ECM fungal communities in temperate oak-hornbeam forests. The results indicate that silvicultural treatments have significant impact on the diversity of ECM fungi and, to some extent, reflect the level of disturbance of the various treatments, confirming our Hypotheses I. For example, differences in alpha diversity of ECM fungi tend to be greatest between control and clear-cut plots, representing the least and most intense disturbance. Most phylogenetic lineages tended to show slightly higher richness in stands with relatively high canopy cover, in intact (C and R) or moderately disturbed (P) plots, and showed decreased richness in CC plots, which suggests that ECM fungi tend to respond negatively to silvicultural treatments that affect the abundance of their host trees. This is in agreement with the findings of the Tedersoo et al. (2014), who found that, at the global scale, alpha diversity of ECM fungi show a strong positive correlation with host abundance.

We also found significant compositional turnover of ECM fungal communities among silvicultural treatments, which confirms our Hypothesis II. In most cases, compositional differences were proportional to the severity of disturbance, e.g., clear-cutting, and to a lesser extent gap-cutting, resulted in the highest species turnover compared to the control, while preparation-cutting and retention tree group only differed slightly from the control. We observed relationships with fungal community turnover and changes in abiotic variables, such as air temperature, relative humidity, and soil water content, suggesting that some effects of the silvicultural practices on fungal communities are mediated through microclimate. These patterns are in agreement with reported changes in abiotic variables in response to these silvicultural treatments, particularly changes in temperature, relative humidity, and light availability (Kovács et al. 2020).

Similarly, fungal community composition correlated with understory plant cover and richness that are also driven by differences in abiotic variables among the treatments (Tinya et al. 2019). Communities in these plots experienced larger fluctuations in temperature and relative humidity due to increased insolation and air movement caused by complete canopy removal. These findings are consistent with Zhang *et al*., (2024), who demonstrated that retention forestry practices moderate microclimate by maintaining canopy cover, while clear-cut areas are subject to higher air temperatures and vapor pressure deficits. The important role of microclimate in shaping the composition of ECM fungal communities has been documented also in tropical forests by Boukary *et al*., (2024) and in Iberian pine forests by Castaño *et al*., (2018). However, it is important to note that while G plots continued to have the “forest microclimate” (Kovács et al. 2020), they tended to differ in composition from the control and were somewhat similar to CC plots, particularly in the 2020 samples. This suggests that gap size is important and that 20-m gaps used in this study likely represent a significant disturbance to ECM fungi, possibly because of the likely decrease in the abundance of available host tree roots in the middle of the plots, i.e. ca.10 m from the forest edge, where the samples were collected. Although not specifically tested in this manuscript, host root abundance is known to be an important factor for the survival of ECM fungi than microclimate, as also reported by Centenaro *et al*. (2024) in Iberian pine forests.

*Amanita* species were mainly found in control and retention tree plots, where the canopy was left intact and, thus, access to host roots was not disturbed, even though the microclimate in retention tree patches is known to be substantially drier than that of unaltered forests (Kovács et al. 2020). Our findings are in agreement with those of Malysheva et al. (2016), who reported greater species richness of *Amanita* in middle-aged to old forests. Similarly, Cullings *et al*. (2005) reported similar reduction in *Cenococcum* abundance in defoliated pine stands due to disrupted host continuity. The ability of *Cenococcum* species to survive in drier forests may be attributed to its melanized sclerotia, specialized structures that confer desiccation tolerance and enable persistence under suboptimal soil moisture conditions (Fogel & Hunt, 1979, 1983; Wang *et al*., 2025). These traits are comparable to those of other disturbance-adapted taxa, such as *Hydnellum peckii*, which survives periods of unfavorable conditions in the form resistant asexual propagules (chlamydospores) (Fernández-Toirán & Águeda, 2007).

The /inocybe lineage showed highest richness in the R plots that generally represent the driest habitat of all treatments in this experiment. This is agreement with the study of Geml et al. (2022) on ECM fungal diversity and composition in several undisturbed Pannonian forest types, who found that, along a temperature and soil moisture gradients, ASV richness of *Inocybe* was the highest in dry oak forests. The relatively high richness observed in several genera in P plots indicate their resilience to moderate canopy alterations, in this case 30% thinning of the dominant canopy and complete removal of the secondary canopy. Also, it reveals that oaks may be the primary host for most of these fungi, because all hornbeam trees were removed from the P plots.

Richness of the /russula-lactarius lineage varied across treatments, with the highest values typically found in control and preparation plots and the lowest in clear-cut areas. Intermediate richness was observed in gap and retention treatments. Their diminished presence in disturbed plots aligns with previous findings reporting lower richness near forest edges and areas affected by disturbance that affect canopy and/or soil structure (Hagerman *et al*., 1999; Kranabetter *et al*., 2013). Furthermore, some *Russula* species are known to persist mainly in old-growth or structurally mature stands, likely due to their reliance on nutrient-mobilizing enzymes that become less effective when root biomass decreases or litter inputs are altered (Rianhard *et al*., 2025b; Twieg *et al*., 2007).

The /tomentella-thelephora clade showed sensitivity to disturbance, with richness notably reduced following clear-cutting but remaining relatively higher in retention and preparation treatments. This pattern indicates that most *Tomentella* species may prefer environments where forest structure and microclimatic conditions are less altered. Similar findings have been reported in other forest ecosystems, where *Tomentella* species increase in abundance as forests mature and disturbance declines (Gao *et al*., 2015). Because compositional changes in /tomentella-thelephora communities relative to the control tended to increase with silvicultural treatment intensity (Figure 4), this lineage may be particularly suitable for bioindicator purposes.

When considering indicator species, faithful and diagnostic to a treatment type, we found that *Amanita citrina, Byssocorticium atrovirens, Inocybe asterospora*, and *Russula grata* were characteristic of the untreated control plots, indicating that these species are sensitive to disturbance. Interestingly, despite representing one extreme of the disturbance gradient that is also visible in NMDS plot of the whole ECM community (Figure 2), the only indicator of the CC treatment was a *Laccaria* species. This genus is known for its ruderal strategy and generally thrives in disturbed forests and urban parks (Hughes *et al*. 2020, Geml, pers. obs.). Conversely, P treatment had the highest number of indicators, suggesting that intermediate disturbance, in this case the thinning of the canopy, provides a niche for the establishment of forest ECM fungi that would otherwise be outcompeted in undisturbed forests. The positive effect of intermediate disturbance on plant communities is well known and examples from fungal studies are gradually accummulating as well (e.g., Adamo et al. 2021).

Niche theory suggests that abiotic factors, such as soil pH, moisture, and temperature, as well as biotic factors, including host abundance and root availability, influence ECM fungal assemblages by acting as environmental filters (Brundrett & Tedersoo, 2020; Mandolini,*et al*., 2024; Suz *et al*., 2014). Along gradients of these factors, there is a spatially heterogenous selection of species based on their physiological tolerances, resource acquisition traits, life strategies (e.g., ruderal, stress-tolerant or competitor), and overall fitness, leading to variation in community composition across habitats and management regimes, as discussed above. In addition, the weaker, but significant compositional differences between the two sampling years indicate a certain level of temporally heterogenous selection that likely is driven partly by interannual differences precipitation. Such annual variations are known to affect fungal growth, reproduction, and host interactions, influencing diversity and functional traits (Fernandez *et al*., 2023; Nuland *et al*., 2024; Sachsenmaier *et al*., 2024). Together, these perspectives provide a basis for understanding the spatial and temporal dynamics of ECM fungal communities in temperate forests (Suz *et al*., 2014; Wilgan & Leski, 2022). It is important to note that differences in richness and compositional patterns of ECM fungi observed in different silvicultural treatments may partly be caused by differences in phylogenetically conserved physiological constraints and functional traits among genus-level evolutionary lineages. In addition species-level differences in habitat preference among congeneric species suggest that there are important ecological differences among congeneric species that drive the dynamics of their competitive interactions along the environmental gradients caused by the silvicultural treatments.

### Conclusions and implications for forest management

Our study shows ECM fungi are sensitive to disturbance and respond strongly to forest management practices. Because forest trees rely on mycorrhizal fungi obtaining water and nutrients, changes in ECM communities expected to have an effect on tree fitness, growth rate and resilience to environmental stressors, e.g., drought stress and pest outbreak. influences belowground biodiversity and ecosystem processes. Forestry practices that maintain canopy continuity, such as retention and low-impact preparation cuts, help preserve disturbance-sensitive fungi like *Amanita citrina, Russula grata*, and *Byssocorticium atrovirens*. These fungi depend on stable microclimates and continuous host-root networks, which are often disrupted by intensive treatments such as clear-cutting. Canopy cover moderates microclimatic extremes in temperature and humidity, supporting moisture-dependent fungi involved in soil nutrient cycling and forest resilience. In clear-cut areas, higher vapor pressure deficits and temperatures favor disturbance-tolerant fungi like *Laccaria trichodermophora*, but community complexity and late-successional species decline. This shift may affect ecosystem functions provided by diverse fungal communities.

Variation in fungal richness and composition in gap-cutting and retention treatments indicates that moderate disturbance can create diverse microsites that support both stress-tolerant and sensitive fungi. The size and arrangement of retained canopy patches likely affect their ability to serve as refuges and sources for recolonization. Considering climate variability that alters temperature and moisture, forest management aimed at maintaining canopy stability and buffering microclimate could support ectomycorrhizal diversity. Including fungal biodiversity in silvicultural planning may improve conservation of these fungi and sustain ecosystem services such as nutrient uptake, soil stabilization, and tree health.

### Future research directions

This study highlights the influence of silvicultural treatments on ECM fungal communities, mediated in part by treatment-driven changes in microclimate and understory vegetation. however, several aspects warrant further investigation. Long-term studies monitoring fungal community composition for several years during forest regeneraton following disturbance could provide more detailed understanding of recovery trajectories and community resilience. Molecular and genomic analyses focused on key fungal clades, such as *Cenococcum*, may help clarify the genetic factors related to stress tolerance and host associations *in situ*. Such insights could contribute to identifying mechanisms of fungal adaptation to abiotic stress, such as decreased soil moisture. In addition, further research examining the influence of canopy gap size and shape on the compositional dynamics of the ECM fungal community will be valuable. Combining fungal community data with measurements of soil properties and forest regeneration will elucidate interactions that affect belowground biodiversity and functioning as well as forest regrowth patterns and will inform stakeholders on the possible consequences and community trajectories resulting from different management practices.

## Acknowledgements

The Pilis Forestry Systems Experiment and related research on were funded by the Hungarian National Research, Development and Innovation Office (NKFIH) PD146325 to B. Kovács, FK145840 to F. Tinya, K 111887 and K143270 to P. Ódor and the Interreg VI-A Hungary-Slovakia Programme (HUSK/2302/1.2/168). The forest microbiome work was funded by an NKFIH OTKA K139387 and the Lendület Program (no 96049) of the Hungarian Academy of Sciences and the Hungarian Research Network awarded to J. Geml. K.O. Ododa is grateful for the support of the Stipendium Hungaricum scholarship program. The authors thank the Pilis Park Forestry Company for their continued logistical support and collaboration.

